# A large-scale chemical-genetic strategy to design antimicrobial combination chemotherapy for *Mycobacterium tuberculosis*

**DOI:** 10.1101/772459

**Authors:** Eachan O. Johnson, Emma Office, Tomohiko Kawate, Marek Orzechowski, Deborah T. Hung

## Abstract

The efficacies of all antibiotics against tuberculosis are eventually eroded by resistance. New strategies to discover drugs or drug combinations with higher barriers to resistance are needed. Previously, we reported the application of a large-scale chemical-genetic interaction screening strategy called PROSPECT to the discovery of new *Mycobacterium tuberculosis* inhibitors, which resulted in identification of the small molecule BRD-8000, an inhibitor of a novel target, EfpA. Leveraging the chemical genetic interaction profile of BRD-8000, we identified BRD-9327, another, structurally distinct small molecule EfpA inhibitor. We show that the two compounds are synergistic and display collateral sensitivity because of their distinct modes of action and resistance mechanisms. High-level resistance to one increases the sensitivity to and reduces the emergence of resistance to the other. Thus, the combination of BRD-9327 and BRD-8000 represents a proof-of-concept for the novel strategy of leveraging chemical-genetics in the design of antimicrobial combination chemotherapy in which mutual collateral sensitivity is exploited.

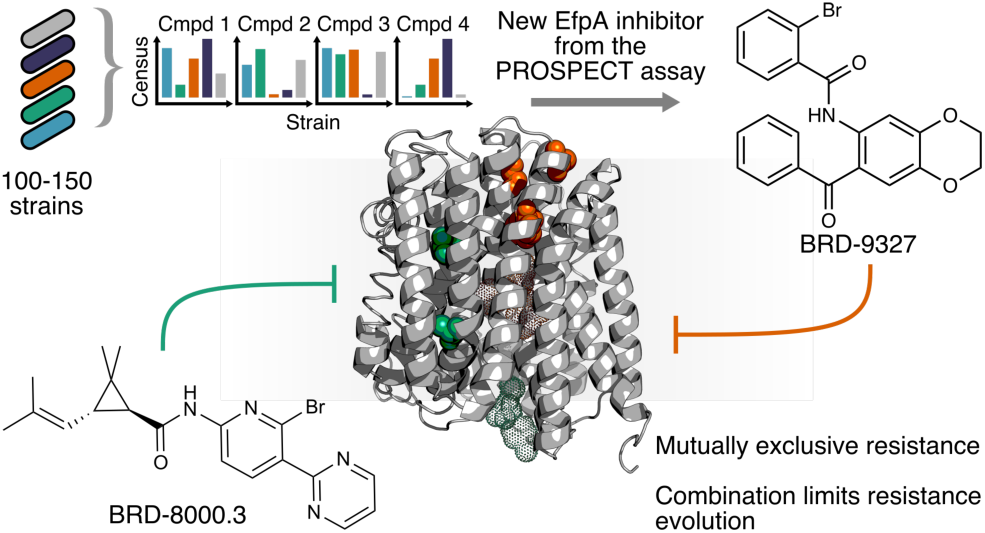

---

Diseases caused by mycobacteria are a significant public health burden, with *Mycobacterium tuberculosis* (Mtb) in particular causing >1.6 million deaths from tuberculosis (TB) annually^1^. The standard of care for drug-susceptible TB is a six-month regimen based on rifampin, isoniazid, pyrazinamide, and ethambutol, but increasing incidence of multi-drug resistant (MDR) TB^1^ is forcing deployment of less effective but longer, more expensive, and more toxic regimens, although improved regimens are in development^2^. With antimycobacterial discovery and development struggling to fill the gaps created by emerging resistance, there is an unmet need for new drugs against TB.

New strategies to discover drugs or drug combinations with higher barriers to resistance are needed. While combination therapy has been the major underlying principle to evade resistance evolution, informed decisions on the best combinations, taking into account the interactions of individual compounds and their resistance mechanisms, has to date been lacking. Here we propose leveraging large-scale chemical interaction studies to identify compound sets that inhibit the same target, thereby enabling the discovery of pairs of compounds that exhibit collateral sensitivity. Collateral sensitivity, that is resistance to a compound that confers hypersensitivity to another, results in a combination whose resistance barrier is higher than two non-interacting compounds.

Previously, we reported a sequencing-based, large-scale chemical-genetic screening strategy, PRimary screening Of Strains to Prioritize Expanded Chemistry and Targets (PROSPECT), which generated chemical genetic interaction profiles (CGIPs) that characterized the fitness of 150 multiplexed, genetically-barcoded hypomorph mutants (strains depleted of individual essential gene products) of Mtb H37Rv in response to ∼50,000 compounds (Figure 1A)^3^. PROSPECT quantifies the fitness changes of genetically barcoded hypomorph strains on compound treatment; the vector of fitness changes, measured as log(fold-change) of abundance of barcodes of a particular hypomorph after treatment with a compound of interest relative to a vehicle control, is known as a CGIP (Figure 1A). Addressing the need for MOA diversity in tackling antimicrobial resistance, PROSPECT can be used to prioritize compounds from primary phenotypic screening data based on their putative MOA, instead of simply their potency. We illustrated PROSPECT’s strengths in the discovery of BRD-8000, an uncompetitive inhibitor of a novel target, EfpA (Rv2846c), an essential efflux pump in Mtb. Though BRD-8000 itself lacked potent activity against wild-type Mtb (MIC_90_ ≥ 50µM), chemical optimization yielded BRD-8000.3, a narrow-spectrum, bactericidal antimycobacterial agent with good wild-type activity (Mtb MIC_90_ = 800 nM, Figure 1B)^3^.

**Figure 1.**
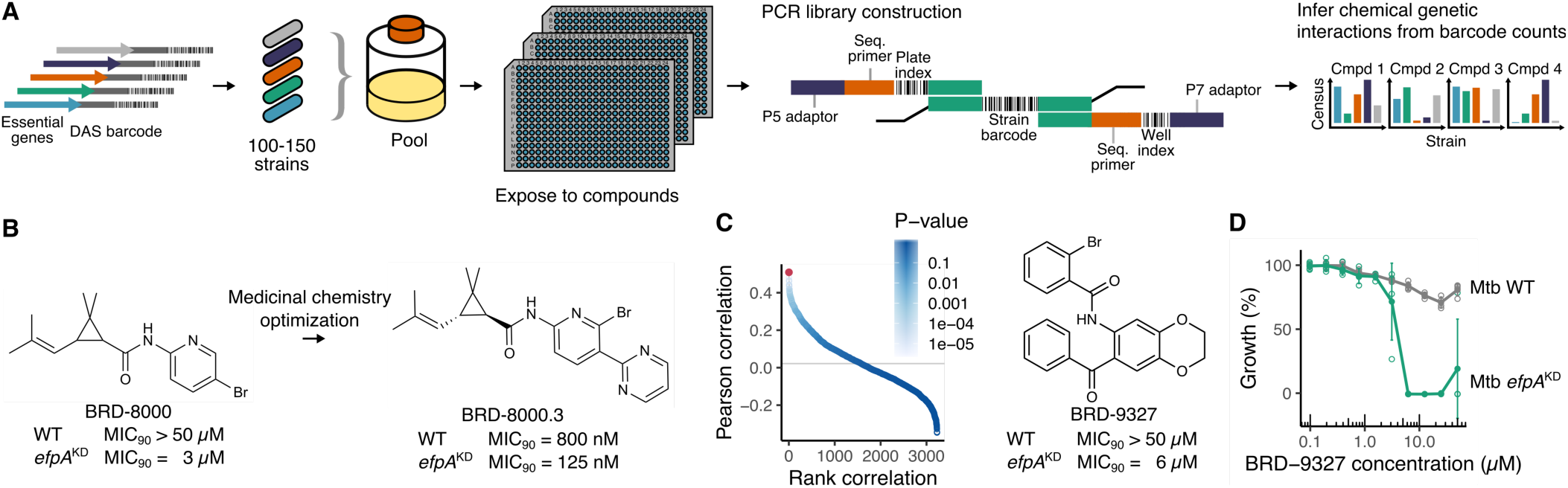
Discovery of a new putative inhibitor of the essential mycobacterial efflux pump, EfpA. (A) Overview of PROSPECT, a sequencing-based, high-throughput chemical-genetic profiling assay. A C-terminal DAS tag, which targets the gene product to degradation by caseinolytic protease (Clp), was integrated at the 3′ end of target genes of interest in the chromosome with concomitant genetic barcoding, which allowed pooling of hypomorph strains. After compound exposure, chromosomal barcodes were PCR amplified, sequenced on the Illumina platform, and analyzed for changes in abundance relative to vehicle controls. For each compound, this generated a vector of strain abundance changes, known as a chemical genetic interaction profile (CGIP). (B) Medicinal chemistry optimization of initial hit BRD-8000, an EfpA inhibitor, yielded BRD-8000.3, a narrow-spectrum antimycobacterial with good wild-type activity. (C) Ranked Pearson correlation of CGIPs with the BRD-8000 CGIP. Each point represents a compound’s CGIP correlation; blue shading indicates P-value under a permutation test (*n* = 10,000). Since BRD-8000 had been validated as an EfpA inhibitor, its CGIP could be used as a reference to discover further EfpA inhibitors. The CGIP of BRD-9327 (highlighted in red) had the highest correlation with the CGIP of BRD-8000. (D) Broth microdilution assay of BRD-9327 against wild-type Mtb and its EfpA hypomorph (Mtb *efpA*^KD^); open circles show individual replicates (*n* = 4), filled circles indicate the mean, and error bars show the 95% confidence interval. BRD-9327 showed very little activity against wild-type Mtb, although the EfpA hypomorph was hypersensitive.

A fundamental strength of PROSPECT is its generation of a large panel of chemical-genetic interactions (7.5 million in the previously reported screen^3^) that can be iteratively and retrospectively mined for new interactions of interest. For example, upon validation of a new a novel inhibitor’s mechanism of action (MOA), its CGIP can be used as a reference for the subsequent discovery of additional scaffolds that work by inhibiting the same target. Taking this approach, we used the CGIP of BRD-8000 to retrospectively identify and prioritize additional putative EfpA inhibitors from the same primary screening data based on their CGIP correlation with BRD-8000’s CGIP (Figure 1C). The chemically distinct molecule BRD-9327 emerged as another possible EfpA inhibitor.

Here, we demonstrate discovery acceleration afforded by PROSPECT and proof-of-concept for a novel strategy which leverages chemical-genetics in the design of compound combinations which inhibit the same target through different mechanisms. We show that BRD-9327 is indeed an uncompetitive inhibitor of EfpA, synergistic with BRD-8000 in both efflux inhibition and mycobacterial growth inhibition. Interestingly, mutations conferring high-level resistance to either of the two compounds, despite only arising in *efpA*, are mutually exclusive and do not confer cross-resistance; in fact, high-level resistance mutations for either compound can cause hypersensitivity to the other compound, thereby lowering the spontaneous resistance frequency to BRD-8000 in a BRD-9327-resistant background. Together, these observations point to the compounds having distinct interactions with EfpA and a strategy in which the pair could be utilized together in a resistance-suppressing combination or resistance cycling regimen. The discovery of BRD-9327 and its interaction with BRD-8000 demonstrates discovery acceleration through PROSPECT’s ability to rapidly prioritize new classes of inhibitors and identify combinations inhibiting a single target with distinct mechanisms of action, thus enabling a new strategy for combatting resistance.

EfpA is an attractive antimycobacterial target since its inhibition was bactericidal and its activity is narrow-spectrum (EfpA is only present in the Actinomycetes); we therefore sought to expand the chemical lead space by identifying new chemotypes for EfpA inhibition. Our previous identification and validation of BRD-8000 and BRD-8000.3 as specific EfpA inhibitors^3^ allowed us to leverage their CGIPs as references for EfpA inhibition. We identified new chemotypes that inhibit EfpA by prioritizing additional putative EfpA inhibitors from the original primary screening data based on their CGIP correlation with the CGIP of BRD-8000 (Figure 1C). This strategy yielded the identification of chemically distinct BRD-9327 as another possible EfpA inhibitor^3^. BRD-9327 showed very weak Mtb wild-type activity (> 50µM) but moderate activity against the EfpA hypomorph (6.25 µM, Figure 1D).

To determine if BRD-9327 is a specific inhibitor of the EfpA efflux pump in Mtb, we took advantage of ethidium bromide (EtBr), a known substrate of EfpA, to measure the impact of BRD-9327 on EtBr efflux rates^4^. EtBr is ∼30-fold more fluorescent when intracellular than when extracellular^5^; this property can be leveraged to measure the efflux-mediated decrease in intracellular EtBr concentration over time (Figure 2A). In the presence of varying inhibitor concentrations, we measured intracellular EtBr fluorescence over time at varying initial EtBr concentration. We then globally fit a modified Michaelis-Menten equation (accounting for Fick diffusion as well as efflux) to the data, obtaining best-fit parameter estimates for the kinetic substrate-free inhibition constant (*K*_i_) and substrate-bound inhibition constant (*K*_i_′)^6^ (Figure 2B).

**Figure 2.**
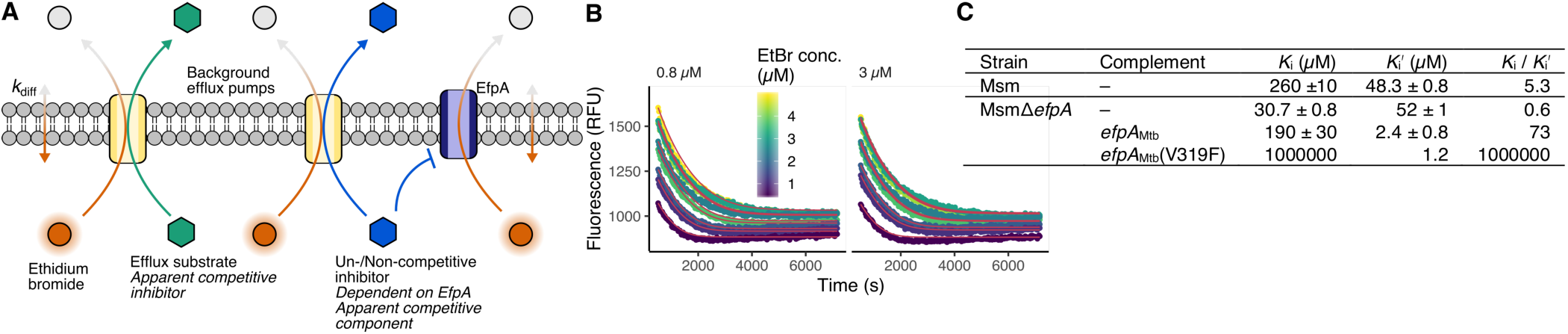
Validating EfpA as the target of BRD-9327 using an EtBr efflux assay. (A) Overview of molecular basis of the EtBr assay for determining kinetic inhibition parameters. When intracellular, EtBr (orange) is ∼30-fold more fluorescent than extracellular; thus, EtBr fluorescence is a proxy for intracellular concentration. In living cells, a compound which is simply a substrate of efflux pumps (green hexagon) will exhibit a competitive mode of EtBr efflux inhibition, since it competes with EtBr for flux through the pumps. However, a compound which has a specific interaction with EfpA (blue hexagon) might also appear to inhibit EtBr efflux competitively, but will exhibit an additional non- or un-competitive modality. In the absence of EfpA, as in a null mutant, this non- or un-competitive modality will be abolished. (B) EtBr fluorescence decay over time (demonstrating varying efflux rates) at ten starting intracellular concentrations and two BRD-9327 concentrations in Msm. Curves corresponding to global best-fit Michaelis-Menten parameter estimates are shown in red. (C) Global best-fit Michaelis-Menten parameter estimates (± standard deviation) of EtBr efflux inhibition by BRD-9327.

We measured EtBr efflux rates in *Mycobacterium smegmatis* MC^2^155 (Msm), a related mycobacterial species, rather than Mtb directly, because Msm’s growth is not affected by BRD-8000 or BRD-9327, presumably because its EfpA homolog (MSMEG_2619) is not essential^7^. We could thus remove the confounding effects of compounds on cellular viability to more cleanly study their direct effect on efflux. However, in addition to EfpA, Msm has a set of other non-essential multi-drug efflux pumps that efflux EtBr. Thus, in order to determine the dependence of the efflux inhibition kinetic parameters on EfpA specifically (Figure 2A), we compared EtBr efflux in a Msm strain containing *efpA* and a strain in which *efpA* had been deleted (Msm*ΔefpA*).

In Msm*ΔefpA*, we found that BRD-9327 is a competitive inhibitor of EtBr efflux by the other multi-drug efflux pumps, with a collective *K*_i_/*K*_i_′ = 0.6 (Figure 2C; *K*_i_/*K*_i_′ < 1 characterizes competitive inhibition). In contrast, BRD-9327 inhibited efflux in the presence of EfpA in wild-type Msm with a *K*_i_/*K*_i_′ = 5.3 (*K*_i_/*K*_i_′ ≥ 1 characterizes non- or un-competitive inhibition; Figure 2C). A mixed or uncompetitive inhibition modality in the presence of EfpA but competitive inhibition in its absence would suggest that while BRD-9327 can be a general efflux substrate of the other efflux pumps, it is a specific, allosteric inhibitor of EtBr efflux by EfpA. Complementation of Msm*ΔefpA* with the Mtb *efpA* homolog showed even more dramatic uncompetitive inhibition (*K*_i_/*K*_i_′ = 100), compared to the wild-type Msm allele, and definitively demonstrated that BRD-9327 is an inhibitor of Mtb EfpA.

We had previously identified a single *efpA* allele in Mtb that confers resistance to BRD-8000 with the V319F amino acid substitution abolishing BRD-8000 binding to mutant EfpA^3^. Interestingly, when we complemented Msm*ΔefpA* with the *efpA(V319F*) allele, while competitive efflux inhibition is observed BRD-8000.3 (due to its activity at the background multi-drug efflux pumps in Msm), we observed uncompetitive efflux inhibition by BRD-9327 (Figure 2C). This uncompetitive inhibition of EfpA(V319F) revealed that BRD-9327 interacts with this mutant EfpA in a manner that must be distinct from BRD-8000’s interaction with EfpA. We therefore tested EtBr efflux inhibition by a combination of BRD-8000.3 and BRD-9327 and found these compounds to be synergistic by excess-over-Bliss (EoB) (Figure S1A).

Having discovered that an allele of Mtb *efpA* that confers resistance to BRD-8000 does not confer biochemical cross-resistance to BRD-9327, we sought to determine if resistance to BRD-9327 would result in cross-resistance to BRD-8000. Because BRD-9327 had not been chemically optimized like the BRD-8000 series to have potent Mtb activity (Mtb MIC_90_ of BRD-9327 ≥ 50 µM), we turned to *Myobacterium marinum* M (Mmar), another related, pathogenic mycobacterial species, that was more sensitive to BRD-9327 (MIC_90_ = 25 µM, Figure 3D).

**Figure 3.**
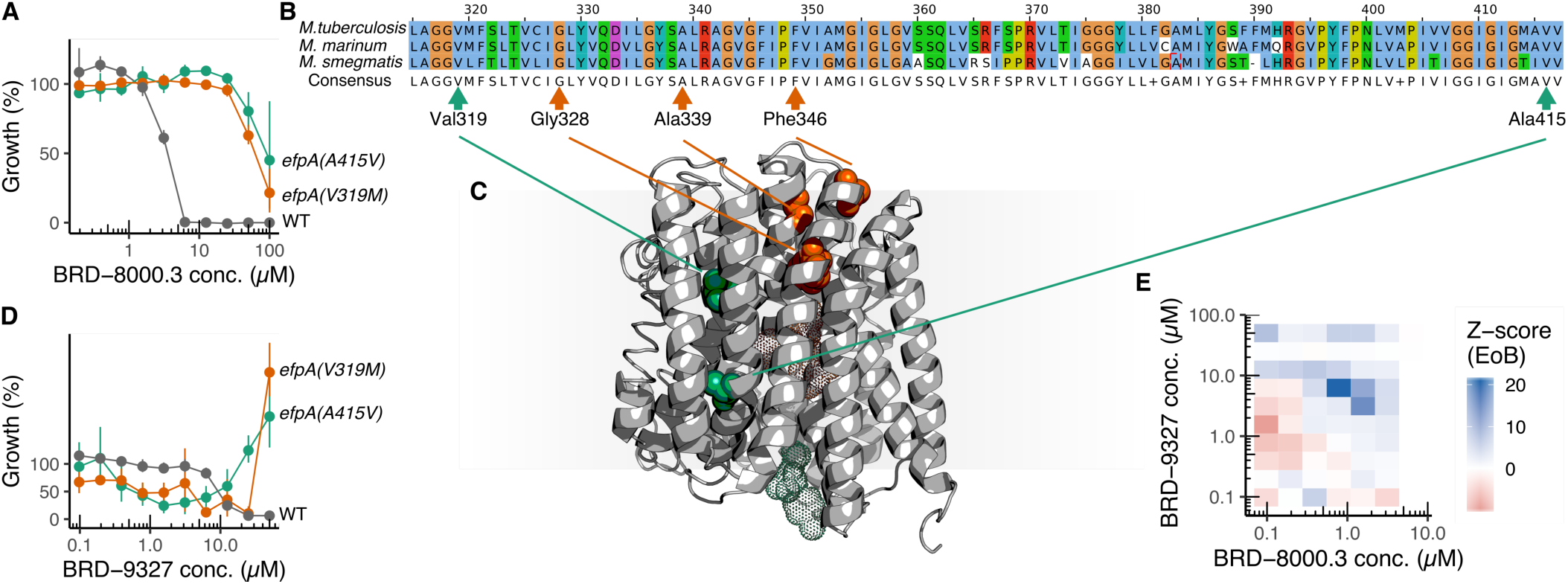
Evolution of Mmar mutants resistant to BRD-8000.3 or BRD-9327. (A) Broth microdilution dose response assay of Mmar and its BRD-8000.3-resistant mutants against BRD-8000.3, demonstrating their high-level resistance to this compound. Filled circles show the mean and and error bars indicate the 95% confidence interval (*n* = 4). (B) Amino acid sequence alignment of highly conserved EfpA in Mtb, Mmar, and Msm, with sites conferring resistance to BRD-8000.3 (green) or BRD-9327 (orange) highlighted. (C) Homology model of EfpA with mutations conferring resistance to BRD-8000.3 (green) or BRD-9327 (orange) highlighted. Mesh outlines show possible binding sites of BRD-8000.3 (green) and BRD-9327 (orange), as determined by docking using AutoDock Vina. (D) Broth microdilution dose response assay of Mmar mutants resistant to BRD-8000.3 against BRD-9327, demonstrating the hypersensitivity of Mmar *efpA(V319M*) and Mmar *efpA(A415V*). Filled circles show the mean and error bars indicate the 95% confidence interval (*n* = 4). (E) Excess-over-Bliss (EoB) of Mmar growth inhibition at varying combined concentrations of BRD-9327 and BRD-8000.3, demonstrating synergy between the two EfpA inhibitors.

We first re-generated BRD-8000.3-resistant mutants in Mmar to provide a baseline comparison of BRD-8000-resistance conferring mutations in Mtb and Mmar. We plated exponentially growing bacteria on agar containing BRD-8000.3 at 2×, 4×, and 8× the broth microdilution MIC_90_ (6.25 µM in Mmar; Figure 3A) to obtain a resistance at a frequency of ∼4 ×10^−8^, confirmed shifts in the broth microdilution MIC_90_ of selected colonies, and performed whole genome sequencing (WGS) of resistant clones on the Illumina MiSeq or HiSeq platform. Whereas we had only observed a single resistance-conferring variant in Mtb (V319F)^3^, we isolated two different Mmar resistant mutants both containing alterations in Mmar *efpA*, V319M and A415V (Figure 3B) which conferred a >16-fold increase in MIC_90_ (Figure 3A). Although there is no high-resolution structure of EfpA, a homology model constructed with I-TASSER^8^ suggested that Val_319_ and Ala_415_ are on neighboring α-helices, and that these mutations could implement the same resistance mechanism (Figure 3C). Consistent with our finding that the *efpA(V319F)* allele of Mtb did not confer functional, biochemical cross-resistance to BRD-9327, BRD-8000.3 resistant mutants of Mmar did not have resistance to BRD-9327. In fact, surprisingly, Mmar *efpA(V319M)* was four-fold more sensitive than wild-type, with MIC_90_ of 6.25 µM for the mutant compared to 25 µM for wild-type Mmar (Figure 3D); although MIC_90_ of Mmar *efpA(A415V)* was > 50µM, it showed IC_50_ of 800 nM. Interestingly, although both BRD-8000 resistant mutants’ growth was inhibited by BRD-9327 concentrations below 25 µM, these strains showed unrestricted growth at BRD-9327 concentrations above 25 µM.

We next sought to identify Mmar *efpA* alleles that confer resistance to BRD-9327. While BRD-9327 is more potent against Mmar than Mtb, its corresponding MIC_90_ is nevertheless too high to allow straightforward selection. Instead, inspired by the efflux synergy of BRD-8000.3 with BRD-9327, we performed a checkerboard assay for growth inhibition of Mmar by the two compounds in combination, and found that they were synergistic by EoB (Figure 3E, Figure S1B). We therefore selected for mutants on agar containing 50 µM BRD-9327 supplemented with 3 µM BRD-8000.3. Since colonies that grew on this combination could escape selection pressure by evolving resistance to either compound, we picked and screened 21 colonies for resistance to each compound individually using a broth microdilution assay. WGS revealed *efpA* variants G328C, G328D, A339T and F346L, which conferred high-level resistance to BRD-9327 but not BRD-8000.3 (Figure 3A). The same homology model of EfpA suggested that these mutated amino acids appeared to reside on neighboring α-helices, again indicating that they could implement the same resistance mechanism (Figure 3C). We identified an additional mutation resulting in a L108Q substitution in *mmar_1007*, the homolog of *Rv0678*, a transcriptional regulator of multidrug efflux pump MmpL5 in Mtb^9, 10^ (Figure S2A) which conferred low-level resistance to both BRD-9327 and BRD-8000.3 (Figure 4A-B), as well as clofazimine (Figure S2B), by increasing expression of MmpL5 and thus efflux of BRD-8000.3 and BRD-9327 (Figure S2C).

**Figure 4.**
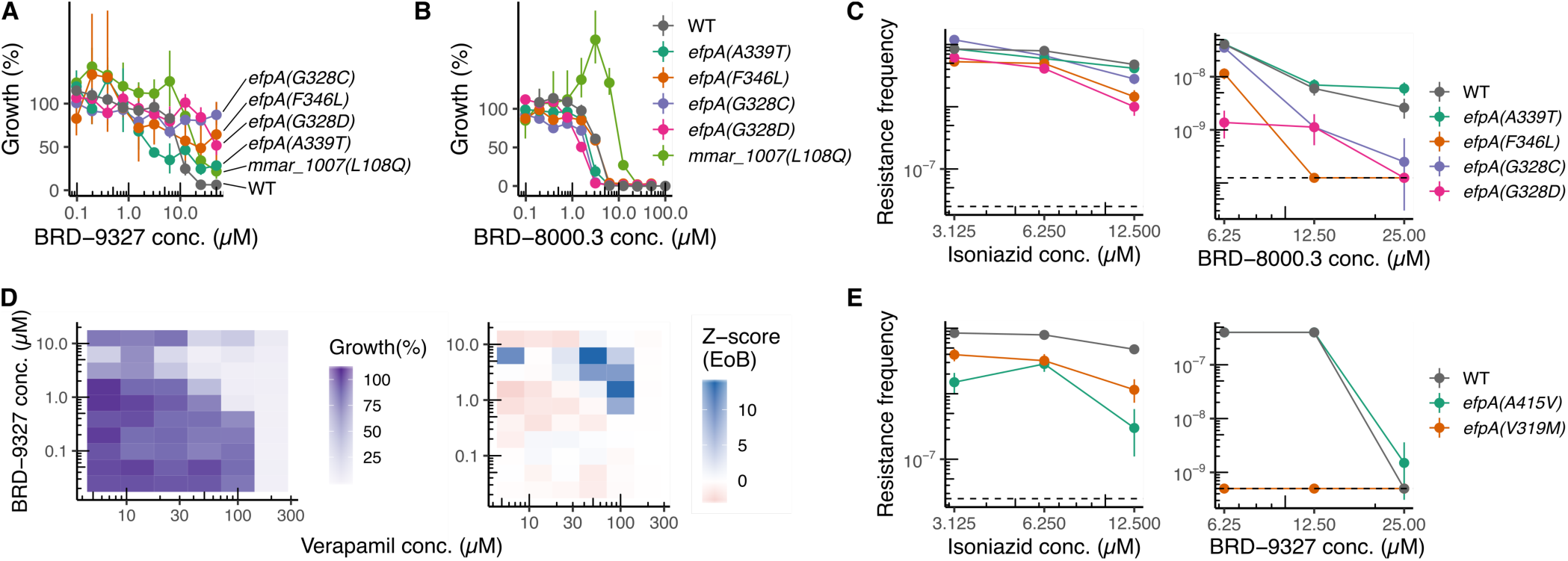
Resistance to BRD-9327 lowers resistance frequency to BRD-8000.3. (A) Broth microdilution dose response assay of Mmar and its BRD-9327-resistant mutants against BRD-9327, demonstrating the high-level resistance of *efpA* mutants, and low-level resistance of the *mmar_1007* mutant. Filled circles show the mean and and error bars indicate the 95% confidence interval (*n* = 4). (B) Broth microdilution dose response assay of Mmar mutants resistant to BRD-9327 against BRD-8000.3, demonstrating the hypersensitivity of Mmar *efpA(G328D)* and Mmar *efpA(A339T)*. Filled circles show the mean and error bars indicate the 95% confidence interval (*n* = 4). (C) Frequency of wild-type or BRD-9327-resistant mutant colonies growing on agar containing 2×, 4×, or 8× MIC_90_ of INH (left) or BRD-8000.3 (right). Filled circles show the mean and error bars indicate the 95% confidence interval (*n* = 4). The dashed line indicates the limit of detection. (D) Growth inhibition from broth microdilution assay of Mmar (left) and the calculated excess-over-Bliss (EoB, right) at varying combined concentrations of BRD-9327 and verapamil, demonstrating synergy between the two compounds. Broth microdilution dose response assay of wild-type Mmar against BRD-9327, verapamil, or the synergistic combination of BRD-9327 with 5 *μ*M verapamil. (E) Frequency of wild-type or BRD-8000.3 resistant mutant colonies growing on agar containing 2×, 4×, or 8× MIC90 of INH (left) or BRD-9327 supplemented with verapamil (right). Filled circles show the mean and error bars indicate the 95% confidence interval (*n* = 4). The dashed line indicates the limit of detection.

In parallel to the mutants resistant to BRD-8000 but hypersensitive to BRD-9327, the resistant mutants of BRD-9327 containing different *efpA* alleles did not exhibit cross-resistance to BRD-8000, and instead, some were hypersensitive to BRD-8000.3. The *efpA(G328C), efpA(G328D)*, and *efpA(A339T)* mutants showed a two-fold decrease in MIC_90_ for BRD-8000.3, while the other mutants with high-level BRD-9327 resistance were not resistant to BRD-8000.3 (Figure 4B). The unique interactions of the two EfpA inhibitors with EfpA, as revealed by their mutual collateral sensitivity pointed to each having a narrow, target-specific resistance space, with mutations disrupting interactions with one compound exacerbating interactions to the other.

Given the mutual collateral sensitivity in the interaction of the two EfpA inhibitors, we speculated that these compounds could be used in a strategy to prevent emergence of high-level resistance. To test this idea, we compared the resistance frequencies for BRD-8000.3 at 12.5µM, 25µM, and 50µM in wild-type Mmar with the those in the Mmar mutants already resistant to BRD-9327. At 12.5 µM BRD-8000.3, while the resistance frequency of Mmar *efpA(F346L)* was 10^−8^, a four-fold decrease compared to wild-type Mmar, the resistance frequency of Mmar *efpA(G328D)* was 2 × 10^−9^, a 20-fold decrease (Figure 4C). Whereas the wild-type resistance frequencies were 6 × 10^−9^ and 2 × 10^−9^ for 25 µM and 50 µM BRD-8000.3, no colonies could be recovered at all for *efpA(F346L)* on 25 µM BRD-8000.3 or higher, nor *efpA(G328D)* on 50µM BRD-8000.3, indicating that BRD-9327 resistance lowers the probability of evolving BRD-8000 resistance (Figure 4C). The *efpA* mutant strains do not have an intrinsically higher mutation rate as the resistance frequencies for isoniazid were identical (3 × 10^−6^).

When we sought to perform the converse experiment to compare the resistance rates for BRD-9327 in wild-type Mmar with the rates in the BRD-8000 resistant Mmar *efpA*_V319F_ mutant, using verapamil as a synergistic potentiator of BRD-9327 to lower its MIC_90_ to permit resistance selection in Mmar (Figure 4D), we again identified a barrier to resistance generation, now for evolving BRD-9327 resistance in a BRD-8000-resistant background. While wild-type Mmar showed unrestricted growth on 6.25 µM and 12.5µM BRD-9327 in the presence of 3 µM verapamil, and the resistance frequency for Mmar *efpA(A415V)* was comparable with wild-type Mmar (∼10^−9^ at 25 µM), no BRD-9327-resistant mutants could be isolated at any concentration for Mmar *efpA(V319M)* (Figure 4E).

The power of large-scale chemical-genetics as a primary screening modality, as implemented in PROSPECT, lies in its ability to incorporate putative MOA information into the prioritization of compounds, moving away from selection simply based on potency. After initial identification of an inhibitor of a new antimycobacterial target, EfpA, PROSPECT allowed for rapid target validation and iterative diversification of chemical scaffold space. With the identification of two chemically distinct EfpA inhibitors, BRD-8000 and BRD-9427, interestingly we identified disjoint sets of target mutations conferring high-level resistance to the two scaffolds. Importantly, resistance to either compound mutually inflicts collateral sensitivity to the other, thereby raising the barrier against resistance to the combination.

The combination of BRD-8000.3 and BRD-9327 is a proof-of-principle demonstration of a novel strategy which leverages chemical-genetics in the design of compound combinations restricting resistance space to a single essential gene, while inhibiting a single target by two different modalities in a manner that makes high-level resistance mutually exclusive. Their unique synergistic interaction illustrates the strategy for combining or cycling therapeutics, with the ability to increase the barriers to drug resistance even in the pursuit of a single target. The use of combination therapy is a critical characteristic of antimycobacterial drug regimens to tackle inevitable resistance evolution to any single agent, which has resulted in the current drug resistance crisis; the identification of rationally designed drug combinations or targets that manipulate the barrier to resistance evolution will be invaluable. This work identifies EfpA as one such valuable target because of its ability to be inhibited by BRD-8000 and BRD-9327 by mutually exclusive mechanisms. Whether EfpA is singularly unique, one of a small number of targets that are amenable to this strategy, or represents a common theme to be more broadly exploited remains to be seen. Nevertheless, this work demonstrates that EfpA is an important and valuable target that can be exploited in this way. Importantly, the ability of PROSPECT to rapidly expand the diversity of scaffolds hitting a single target, as illustrated for EfpA, will enable the potential discovery of complementary inhibitors with variable mechanisms of action and facilitate greater exploration and expansion of this targeting strategy not only to tackle increasing tuberculosis drug resistance, but also more generally, to tackle other resistant pathogens and diseases such as cancer.

## Author information

### Author contributions

The project was devised by E.O.J. and D.T.H. Experiments were designed by E.O.J. Experiments were carried out and results were analyzed by E.O.J. and E.O. Compounds were synthesized by T.K. Homology modeling and docking was carried out by E.O.J. and M.O. E.O.J. and D.T.H. wrote the manuscript.

## Notes

The authors declare no competing interest.

## Acknowledgments

Funding was provided by the Broad Institute TB Gift Donors and Pershing Square Foundation. The authors declare no competing interests.

## Methods

### Strains

The bacterial strains we used and designated as wild-type were *M. tuberculosis* H37Rv, *M. smegmatis* mc^2^155^11^, and *M. marinum* M. Construction of the *M. smegmatis* Δ*efpA* strain and expression constructs for *M. tuberculosis efpA* and *efpA(V319F)* were described previously^3, 12^.

### Compounds

BRD-8000 and BRD-8000.3 were synthesized and characterized as described previously^3^. BRD-9327 was purchased from ChemBridge (catalog #7025440).

### Efflux assay

Efflux rates were measured as previously described^3^. Briefly, Msm strains were grown in Middlebrook 7H9 medium (M7H9) supplemented with oleic acid, albumin, dextrose, and catalase (OADC; BD) to an OD_600_ of 0.4–0.6. Cultures were then centrifuged for 5 min at 3500 rpm. The pellet was washed once with phosphate buffered saline (PBS) at 37 °C and resuspended in 37 °C PBS to give a final OD_600_ of 0.4. Cultures were split into eight and EtBr was added at a final concentration of 0.2-1.95 µg/mL and bacteria were incubated for 30 min (Msm) at 37 °C. After EtBr treatment, cells were centrifuged for 5 min at 3500 rpm and resuspended in 37 °C PBS to give a final OD_600_ of 0.8. A white 96-well plate (Corning) was prepared with serially diluted compound and 50µL PBS containing 0.8% w/v glucose. 50µL dye-loaded bacteria were added to each well of the plate. Fluorescence was read at 37 °C in a SpectraMax M5 plate reader using 530 nm excitation and 585 nm emission wavelengths for EtBr and was recorded every 30 s for 2 h (Msm).

To infer kinetic parameters, we modeled the rate of fluorescence decay as a modified Michaelis-Menten equation, which included a term for Fick diffusion^13^ between the cytoplasm and extracellular mileu, as previously described^3^. Initial efflux rates to determine synergy were calculated by fitting a spline (function smooth.spline^14^ in R) to each time-course and calculating the first derivative at 480 seconds (to avoid knots in the spline).

### Broth microdilution assays

The minimum inhibitory concentration of compounds was determined in a 96-well plate (Corning), filled with 49 µL of M7H9-OADC, and 1 µL 100× compound DMSO stock. 50 µL exponential-phase bacterial culture diluted to an OD_600_ of 0.005 was added. Plates were incubated at 37 °C in a humidified container for 3 d for Mmar, and 14 d for Mtb. OD_600_ was measured using a SpectraMax M5 plate reader (Molecular Dimensions). Normalized percent outgrowth (NPO) was reported using NPO = (*x*_*i*_ *– µ*_n_) / (*µ*_p_ – *µ*_n_), where *µ*_p_ is the mean positive control value, *µ*_n_ is the mean negative control value, and *x*_*i*_ is the value of compound *i*.

### Checkerboard assays and synergy

A 96-well plate (Corning) was filled with 48 µL of M7H9-OADC, and 1 µL of each 100× compound DMSO stock. 50 µL exponential-phase bacterial culture was diluted to an OD_600_ of 0.005 before being added. Synergy was calculated using excess-over-Bliss, which compares the expectation of independent compound effects to the observed combined effect:

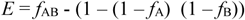

where *E* is excess-over-Bliss, *f*_AB_ is the observed, combined fractional inhibition by the two compounds, and *f*_A_ and *f*_B_ are the observed individual fractional inhibition by each compound. The Z-score of EoB was calculated as *E* / *s*_*E*_, where *s*_*E*_ is the estimated standard deviation of the EoB, calculated by propagating the standard deviations of the underlying growth or efflux rate measurements.

### Evolution of resistant mutants

Mid-exponential growth phase bacterial cultures were pelleted and resuspended at 2 × 10^10^ cfu mL^−1^ in M7H9-OADC. 50 µL (10^9^ cfu) was plated in duplicate on 6 mL M7H10-OADC agar containing 2×, 4× or 8× MIC_90_ of test compound. Plates were incubated at 37 °C in a humidified container. This was repeated on two separate days. At 14 d, agar was checked every 7 d for colonies, which were transferred into 10 mL M7H9-OADC cultures which were grown to mid-exponential phase before testing for resistance in a broth microdilution assay. Resistant mutants were then subjected to whole genome sequencing.

### Whole genome sequencing of Mycobacteria

10 µL of bacterial culture was combined with 10 µL 10% v/v DMSO in a 96-well clear round-bottom plate (Corning). Plates were heat-inactivated at 80 °C for 2 h. Genomic DNA (gDNA) was separated from intact cells and cell debris using AMPure XP (Beckman), eluting in 40 µL MilliQ water. 1.5 µL gDNA was amplified using 6 µM random primers (Invitrogen) and φ29 DNA Polymerase (NEB) 10 µL reaction volume at 30 °C for 24 h.

Amplified gDNA was purified using AMPure XP and subjected to NextEra XT NGS library construction (Illumina) before 150-cycle paired-end sequencing on the Illumina MiSeq platform. Reads were aligned to the CP000854 reference sequence^15^ using the BWA-mem^16^ algorithm and mutations were called using the Genome Analysis Toolkit (GATK)^17^.

### Computational modeling of proteins and ligands

A homology model of EfpA was built using the I-TASSER algorithm^8^, which builds a model from an ensemble of templates, each of which has some sequence homology to a region of the query. For the essential efflux pump, EfpA, I-TASSER used peptide and oligopeptide transporters (PDB 4IKV^18^, 4Q65^19^, 4W6V^20^, 6EI3^21^, 6GS1^22^), human glucose transporter GLUT1 (4PYP^23^), *E. coli* multidrug transporter MdfA^24^ (4ZOW), and *E. coli* organic ion transporter DgoT (6E9N^25^) as templates.

Possible binding sites of BRD-8000.3 and BRD-9327 in the I-TASSER model were calculated using the AutoDock Vina^26^ extension of UCSF Chimera^27^.

