## Supplemental Figures for "A large-scale chemical-genetic strategy to design antimicrobial combination chemotherapy for *Mycobacterium tuberculosis*"

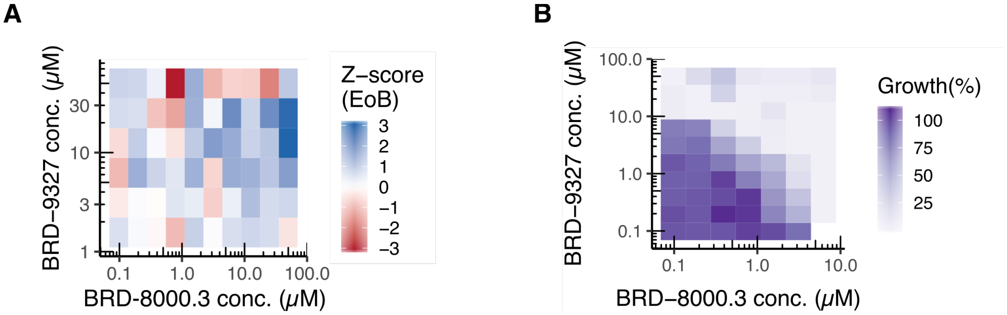


**Figure S1. Synergy between EfpA inhibitors BRD-8000.3 and BRD-9327.**
(A) Excess-over-Bliss (EoB) of initial EtBr efflux rate inhibition in Msm at varying combined concentrations of BRD-9327 and BRD-8000.3, demonstrating synergy between the two EfpA inhibitors.
(B) Growth inhibition from broth microdilution assay of Mmar at varying combined concentrations of BRD-8000.3 and BRD-9327, demonstrating synergy between the two compounds.

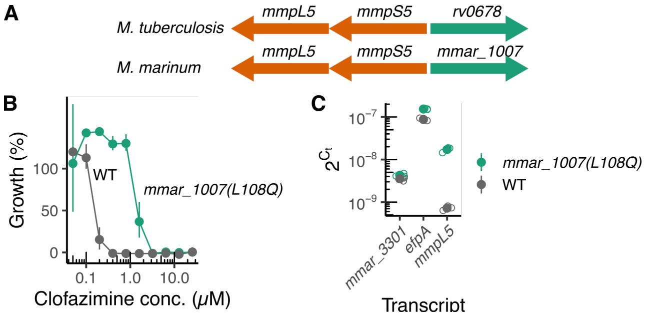


**Figure S2. Low-level cross resistance between BRD-9327 and BRD-8000 is mediated by mutation in the efflux pump MmpL5 regulator *mmar_1007*.**
(A) Chromosomal organization of the MmpL5 pump regulator Rv0679 (Mtb) and MMAR_1007 (Mmar), demonstrating synteny.
(B) Broth microdilution assay of wild-type Mmar and Mmar *mmar_1007(L108Q)* against clofazimine, demonstrating resistance.
(C) Results of a qRT-PCR assay of *mmpL5* and *efpA* transcripts in an *mmar_1007* mutant compared to wild-type Mmar, demonstrating 19-fold overexpression of the multidrug efflux pump MmpL5 in *mmar_1007* mutants.
